# Robust and efficient software for reference-free genomic diversity analysis of GBS data on diploid and polyploid species

**DOI:** 10.1101/2020.11.28.402131

**Authors:** Andrea Parra-Salazar, Jorge Gomez, Daniela Lozano-Arce, Paula H. Reyes-Herrera, Jorge Duitama

**Affiliations:** Department of Systems and Computing Engineering, Universidad de los Andes, Bogotá, Colombia; Corporación Colombiana de Investigacion Agropecuaria (AGROSAVIA), Bogotá, Colombia

## Abstract

Genotype-by-sequencing (GBS) is a widely used cost-effective technique to obtain large numbers of genetic markers from populations. Although a standard reference-based pipeline can be followed to analyze these reads, a reference genome is still not available for a large number of species. Hence, several research groups require reference-free approaches to generate the genetic variability information that can be obtained from a GBS experiment. Unfortunately, tools to perform de-novo analysis of GBS reads are scarce and some of the existing solutions are difficult to operate under different settings generated by the existing GBS protocols. In this manuscript we describe a novel algorithm to perform reference-free variants detection and genotyping from GBS reads. Non-exact searches on a dynamic hash table of consensus sequences allow to perform efficient read clustering and sorting. This algorithm was integrated in the Next Generation Sequencing Experience Platform (NGSEP) to integrate the state-of-the-art variants detector already implemented in this tool. We performed benchmark experiments with three different real populations of plants and animals with different structures and ploidies, and sequenced with different GBS protocols at different read depths. These experiments show that NGSEP has comparable and in some cases better accuracy and always better computational efficiency compared to existing solutions. We expect that this new development will be useful for several research groups conducting population genetic studies in a wide variety of species.

## 1 Introduction

In the last decade, the development and use of high-throughput sequencing (HTS) technologies enabled the production of large volumes of genetic sequence data for almost any form of life. A wide variety of protocols have been developed aiming to maximize the use of HTS data for different applications. Among them, several protocols based on DNA digestion with restriction enzymes and sequencing of the resulting fragments, collectively called Genotype-By-Sequencing (GBS), are being extensively used by a large number of researchers in plant and animal genomics [6]. Some of the most used methods include Restriction Amplification and Digestion (RAD) sequencing [3], single enzyme GBS [10], double enzyme RAD-seq known as ddRAD [25], double enzyme GBS [31], among others. GBS protocols are a cost-effective strategy to discover and genotype thousands of single nucleotide polymorphisms (SNPs) in populations of hundreds or even thousands of individuals [6]. Starting from pioneer studies on three spine stickleback [13], maize [29], and polyploid species such as switch grass [17], these methods have been applied to assess genomic diversity in a large number of plant and animal species [1, 30] with different goals including assessment of genetic diversity in natural populations [14], Genome-Wide Association [19, 2], construction of genetic maps and QTL analysis in controlled populations [12], and genomic prediction [16]. Although whole genome sequencing (WGS) is a viable alternative to obtain complete information of genomic variability, GBS protocols are the preferred choice for species with complex repetitive and polyploid genomes and for controlled and large populations for which most individuals will not be conserved over time.

If a reference genome is available for the species under study, a standard mapping and variants discovery bioinformatic analysis pipeline can be followed to analyze GBS reads [24]. However, this is still not the case for a large number of species. Current biodiversity studies require software tools to perform a reference-free analysis of GBS data. Because GBS reads are naturally organized in non overlapping discrete stacks, standard de-novo assembly packages can not be used to analyze these reads. Different software tools have been developed to perform de-novo analysis of GBS data [4, 11]. Each of these tools implements a different algorithm that performs clustering of reads by similarity, under the assumption that two reads with small differences are sequenced from a single genomic loci and that these differences are due to sequencing errors or real variation within the population. A mock reference is created by obtaining a consensus of the clustered reads to perform variants detection and genotyping. These strategies should be robust in the presence of sequencing errors and other issues occurring at different stages of the GBS protocols [21]. Table 1 summarizes some of the relevant characteristics for the most popular softwares for de-novo analysis of GBS data.

**Table 1:**
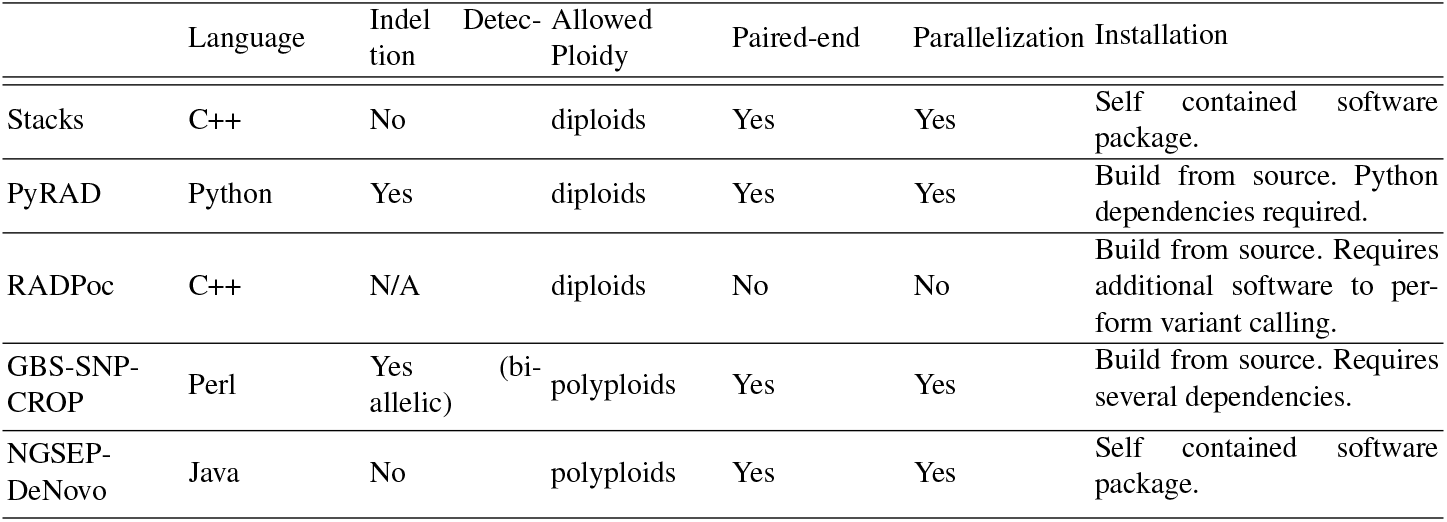
Comparison of the capabilities of available de-novo software tools.

Differences in implementation are found in clustering phase, consensus building and variant calling. RADProc and Stacks rely on an exact k-mer clustering algorithm, followed by a second step to align consensus sequences and group clusters that are likely to come from the same genomic locus. The expectation is that polymorphic sites will produce two clusters, each representing alternative alleles [5]. PyRAD and GBS-SNP-CROP on the other hand employ USEARCH and VSEARCH respectively, which integrates centroid-based clustering algorithms [9, 28]. For consensus calling both PyRAD and GBS-SNP-CROP use multiple alignment. On the other hand, Stacks splits the reads within each cluster in k-mers (default 31) and assemble a contig choosing the path with highest coverage on a DeBruijn graph built from the k-mers. If a reverse contig is successfully assembled, a suffix tree is used to attempt to combine it with the forward contig [27]. Finally, three different approaches are implemented for variant calling. GBS-SNP-CROP calculates the average allele ratio observed across all heterozygous individuals, and a binomial distribution is then used to select variants [22]. PyRAD implements a maximum likelihood equation [18] to estimate average heterozygosity and distinguish variants from sequencing errors [7]. Stacks implements a Bayesian genotype caller developed by Lynch et. al [20]. In all cases, benchmarking is poerformed using simulated data ([22,27,7]). Independent works performed benchmarking for Stacks on real populations [21].

In this manuscript we describe a new algorithm for de-novo analysis of GBS reads. We developed this algorithm to address two issues identified in our experiments with different datasets with current tools for GBS data analysis. First, almost all tools (except for Stacks) lack accuracy in different real datasets, and Stacks can take several days to process large populations. And second, only GBS-SNP-CROP supports genotyping of polyploids but it is not possible to execute it if barcodes are not available for the population. The next sections describe details of this algorithm and benchmark experiments showing that our approach achieves better or comparable performance at lower computational cost compared to current solutions.

## 2 Results

### 2.1 A new algorithm of reference-free analysis of GBS reads

We developed an efficient and robust algorithm for de-novo analysis of Genotype-by-sequencing (GBS) data, which supports different Genotype-By-Sequencing protocols, including ddRAD-seq and SLAF-seq. The algorithm has three main components. Initially, an efficient voting clustering algorithm groups reads considered to be sequenced from the same genomic site. This is done by clustering k-mers within the region of the read that has the highest sequencing quality. This allows for robust clustering considering sequencing errors and possible sequence variation. Then, the algorithm assigns each read to a cluster based on its representative k-mer. Finally, each cluster is internally aligned to generate a consensus sequence and perform variant calling (Fig. 1).

**Fig. 1:**
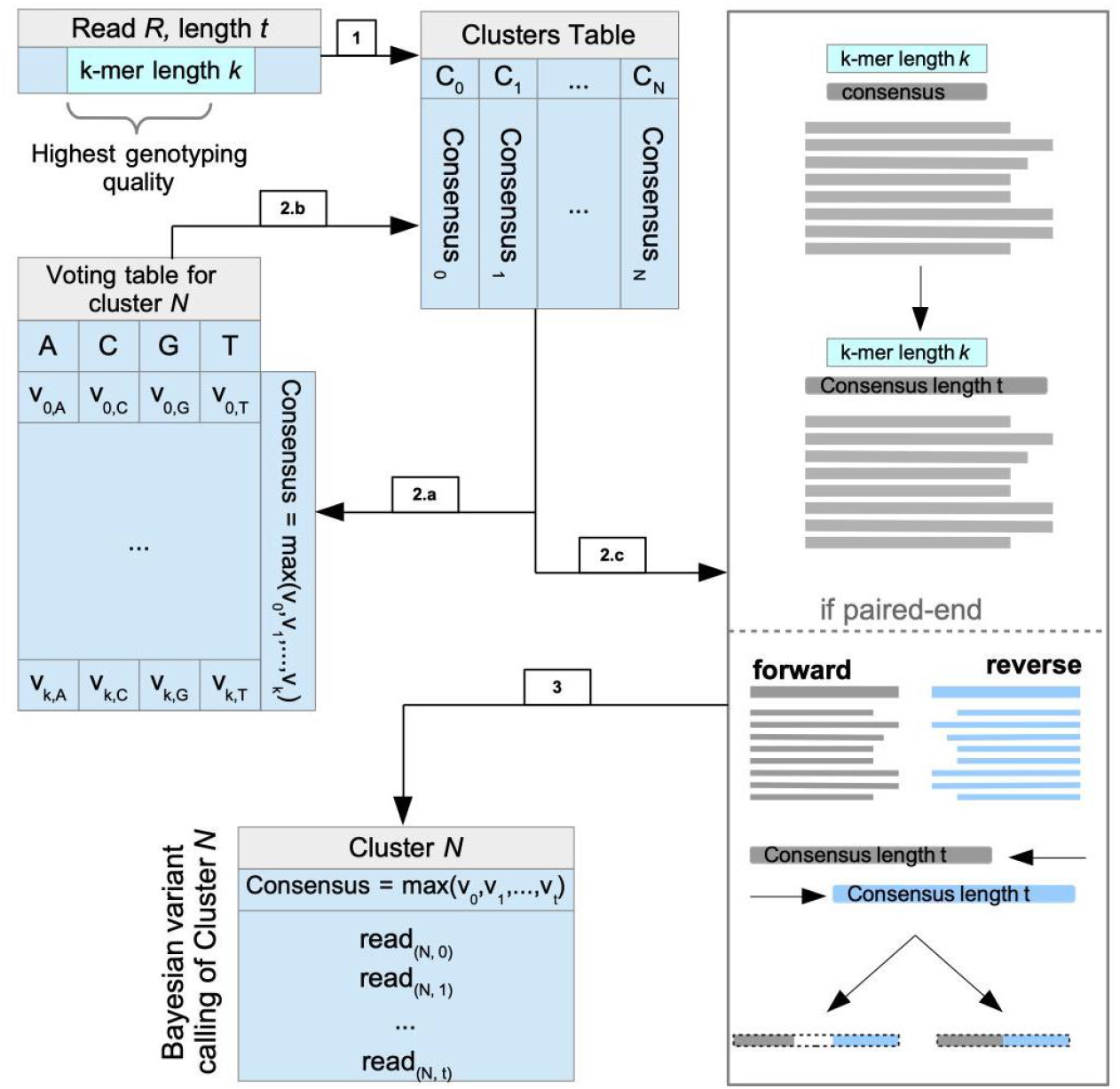
NGSEP-DeNovo algorithm workflow. **1**. Each read will be represented by a k-mer, each cluster is represented by its consensus k-mer. **2.a-b**. If a read is matched with a cluster the voting table for that cluster is updated, or else a new table is created. **2.c**. the consensus is expanded to span the longest read of the cluster. For paired-end reads, consensus sequences are aligned. **3**. Variant calling is performed using consensus sequence as reference.

To perform an efficient identification of clusters, reads are traversed and a representative k-mer is extracted for each read, starting from the sixth position from the 5’ end of the read and with length k (31 by default). The k-mer is not extracted from the first position to avoid covering the portion of the enzyme recognition site still present in the reads. Given a maximum number of clusters M (2 million by default), a counts table of M*k rows and 4 columns is created to count the number of times each base pair appears within each cluster and keep updated the cluster consensus through the first traversal of the reads (Fig. 1.1). Consensus sequences are stored in a separate hash table with a value indicating the start position of the cluster in the counts table. The very first read will create a new cluster for which it will be the consensus. For each read afterwards, the representative k-mer and all its 1-bp neighbors are searched in the consensus hash table to assess if the read should make a new cluster or if it should be added to an existing one. In the former case, the read is added to the cluster, the counts table is updated and the consensus sequence is updated if needed. In the later case, a new cluster is created (Fig. 1.2.a-b.).

Once clusters and consensus sequences are calculated, raw reads are traversed a second time to assign each read to a cluster and to sort the reads of each sample by cluster ID. The representative k-mer of each read is calculated and queried along with its 1-bp neighbors to assign it to its closest cluster. Reads that are not assigned to clusters at this stage are discarded. Sorted reads are dumped in temporary files to keep the process memory tractable.

The temporary files with reads sorted by cluster are traversed in parallel to collect the reads assigned to each cluster and process each cluster in parallel. Reads are aligned to each other using a simple Hamming distance approach, assuming that in GBS data a low percentage of clusters span real indels. A consensus across the entire length of the reads is recalculated by majority vote on the alignment (Fig. 1.2.c). Finally, Variants are called within each cluster using the reference-based variant detection and genotyping algorithm implemented in NGSEP [34] (Fig. 1.3).

For paired-end reads, the clusters are created with the forward reads, using the same approach implemented for single-end reads (Fig. 1.2.c). In the second step, both forward and reverse reads are sorted to keep the read pair information. In the third step, left and right reads are aligned separately and consensus are calculated separately. Then, the consensus sequences of the forward and reverse reads are aligned to determine if they have a significant ungapped overlap. In such case, the two consensus are merged in a single consensus according to the overlap. Otherwise, the reverse complement of the reverse consensus is calculated and attached to the forward consensus leaving a fixed number (5 by default) of unknown sequence character (“N”) in the middle. To perform variant calling, forward reads are left aligned to the merged consensus and the reverse complement of each reverse read is right aligned to the merged consensus.

### 2.2 Benchmarking NGSEP de-novo

We performed bechmarking of the algorithm against GBS de-novo analysis software such as Stacks, RADProc and PyRAD using two different datasets (rice and Sea bass). The third data set was used to evaluate performance in a polyploid species.

The selected datasets have different population structures, GBS protocols, species and ploidies (See methods section 4.2 for details regarding the data). Because in all cases a reference genome was available for the species, we used the variants and genotype calls generated by a reference-based analysis pipeline as gold-standard to evaluate the accuracy of the different software tools (See methods section 4.4 for details regarding accuracy measures).

To match calls generated by reference-based pipelines with calls generated by de-novo pipelines, we developed a *translator,* which aligns consensus sequence of each cluster to a reference genome and translate the variants from the coordinates of the de-novo VCF which are cluster specific (e.g. Cluster 10, position 27) to the coordinates of the reference Genome (e.g. Chromosome 2, position 12,542). This tool not only allowed to validate the different tools with real data sets but also allows to map de-novo variation datasets when a reference genome becomes available for the species under study.

### 2.3 Benchmark with a rice biparental population

The first dataset is a biparental F6 population of rice *(Oryza sativa)* which was one of the first populations genotyped using single reads GBS [32]. Rice is a model species with a high quality reference genome available. On the other hand, this particular library has an average depth of coverage of just 5x, which allows to evaluate the behavior of different software tools on low read depth cases. The algorithm described in the previous section was compared to Stacks, RADproc and PyRad in this dataset.

The first observed impact of low coverage on data quality is the high percentage of missing data reported by all tools, with the exception of NGSEP. (2.A). Although NGSEP called less raw variants compared to the other tools, the raw percentage of missing data was 33%, compared to more than 80% obtained with other tools. Missing data also increases once filtering of genotype calls by raw depth or genotype quality score is applied. The distribution of quality scores for NGSEP genotype calls, both in the de-novo analysis and the reference-based analysis follows a similar path to that of Stacks de-novo (Supplementary figure QSDistRice). However the quality scores of NGSEP are distributed between 0 and 255, whereas the quality scores of Stacks are distributed between 13 and 40. The quality scores of the reference-based analysis implemented in Stacks have the same limits but follow a more conservative distribution.

To perform an accuracy comparison between the different tools, first we followed the same approach used in previous works contrasting the number of segregating variants with at least 100 individuals genotyped against the number of incorrect genotype calls according to the structure of the population [24]. Figure 2.B shows that the de-novo analysis implemented in NGSEP has better accuracy in this dataset than the de-novo analysis implemented in Stacks. The figure shows for comparison the results of the same analysis using reference-based approaches. The de-novo analysis of NGSEP has comparable accuracy with the reference-based approach implemented in GATK and it only falls below the reference-based analysis of NGSEP and Stacks. Even without a filter on minimum quality score, PyRAD and RADproc could genotype less than 1,000 SNPs in 100 individuals, whereas NGSEP and Stacks genotyped about 40,000 SNPs. Based on this result and to make a more direct assessment of genotyping accuracy, we compared genotype calls produced by de-novo and reference approaches. Figure 2.C shows that the genotype calls obtained with the de-novo analysis of NGSEP have better consistency with the reference-based analysis compared to those obtained with Stacks. Whereas NGSEP identified 0.7 million calls consistent with the reference-based approach, Stacks can only identify up to 0.5 million. Assuming that most heterozygous calls are erroneous taking into account that the population has six rounds of inbreeding, Stacks shows a larger number of genotyping errors compared to NGSEP and even compared to the number of non-reference genotype calls in SNPs that could not be matched with SNPs identified by the reference-based approach.

**Fig. 2:**
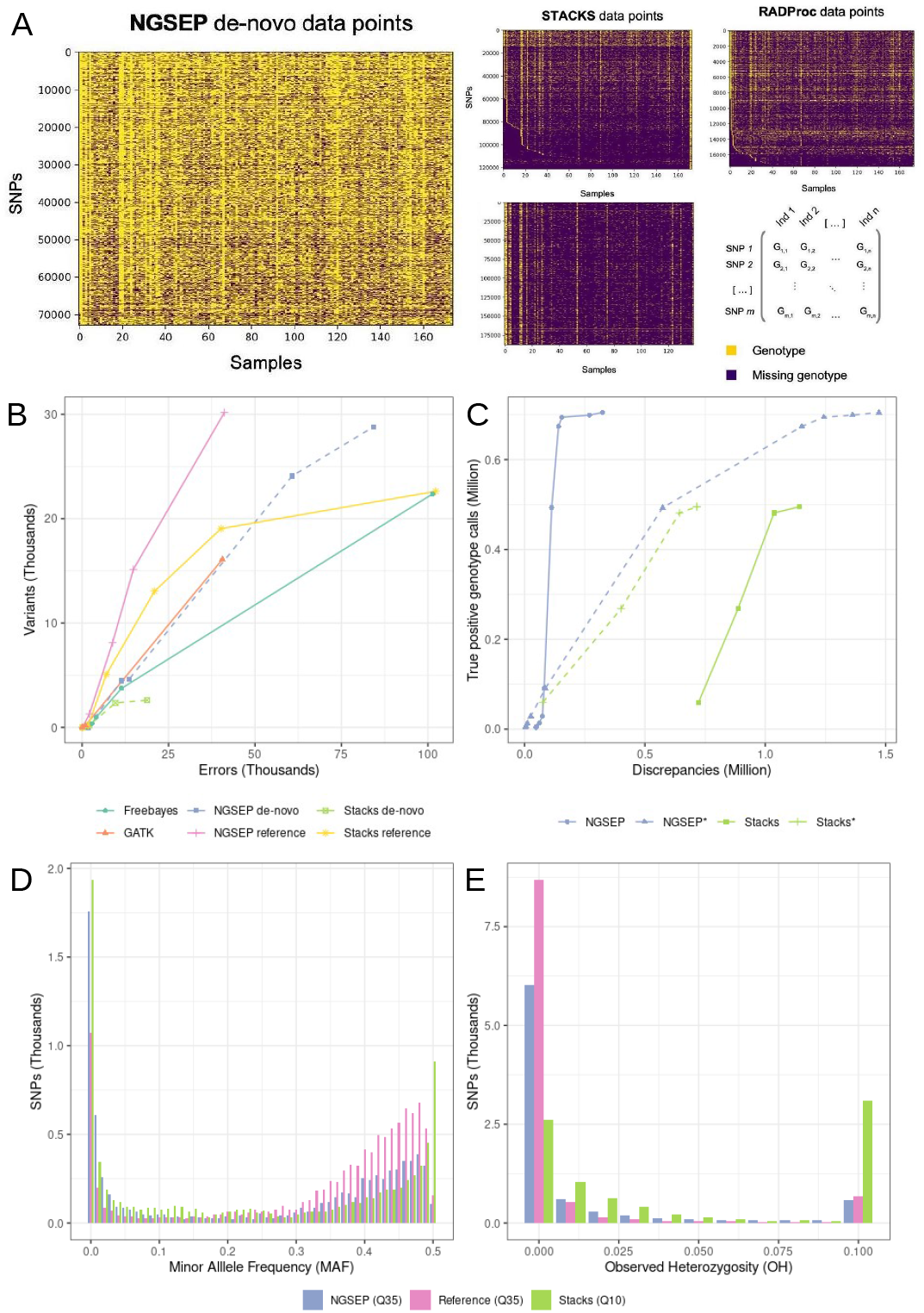
**A.** Overview of raw genotype calls obtained with NGSEP, Stacks, RADProc and PyRAD over the 173 individuals of the rice Azucena x IR64 biparental population. **B.** Number of segregating SNPs genotyped in at least 100 individuals as a function of the number of genotyping errors according to the population structure. De-novo approaches are differentiated with dashed lines. **C.** Number of matching non-reference genotype calls between de-novo and reference-based approaches as a function of two types of discrepancies: genotype differences within matching SNPs (solid lines) and non-reference genotype calls in SNPs identified only with the de-novo approach (NGSEP* and Stacks* dashed lines). **D-E**. Distribution of minor allele frequency (D) and observed heterozygosity (E) for SNPs genotyped with NGSEP and Stacks. The same curves obtained with the reference-based analysis are shown for comparison

Further validation of the results was conducted taking into account the distribution of population statistics expected for a biparental population. As expected, the observed distribution of MAF has a peak close to 0.5 for variants generated with each tool (Fig 2.D). Stacks shows the larger number of SNPs with low MAF, which are likely to be errors. Stacks also shows a larger peak for SNPs with MAF 0.5. However, the distribution of observed heterozygosity (OH) suggests that these SNPs also have a large OH, which is not expected for an F6 population (Fig 2 E). Conversely, SNPs generated by the de-novo approach implemented in NGSEP show the expected distributions of MAF and OH. Compared to the reference-based approach the main differences are a larger number of errors and a lower number of identified SNPs with high MAF.

### 2.4 Sea Bass diversity population

As a second benchmark dataset, we used a Seabass natural population of 146 individuals genotyped with double digest RAD sequencing (ddRAD) [25]. A raw number of 433,158 and 491,957 variants were called using NGSEP and Stacks respectively. Although a higher average read depth per sample translated in a lower missing data rate, especially for Stacks, NGSEP also showed a lower raw missing data rate in this case (Supplementary figure 2). After mapping to a reference genome, an accuracy comparison similar to that performed on the rice population was performed in this case. However, heterozygous calls in this population should be as common as homozygous alternative calls, the analysis was performed separately for heterozygous and homozygous calls. Figure 3.A shows that Stacks calls about 0.2 million matching more datapoints compared to NGSEP with a comparable rate of genotyping discrepancies in matching SNPs. However, Stacks also generates about two times more non-reference genotype calls outside matching SNPs compared to NGSEP (dashed lines). Focusing on genotyping discrepancies in matched sites, whereas for NGSEP 80% of the discrepancies are found in heterozygous calls according to the reference-based analysis, this percentage is close to 40% for Stacks. This suggests that the errors produced by the de-novo analysis implemented in NGSEP are probably due to lack of clustering of reads in some truly heterozygous sites, which can be a consequence of the less efficient read clustering algorithm. Conversely, the errors produced by Stacks seem to be more related to the variant discovery algorithm. Whereas for NGSEP, less than 8% of the genotype discrepancies are homozygous calls to different alleles, this percentage is 25% for Stacks.

**Fig. 3:**
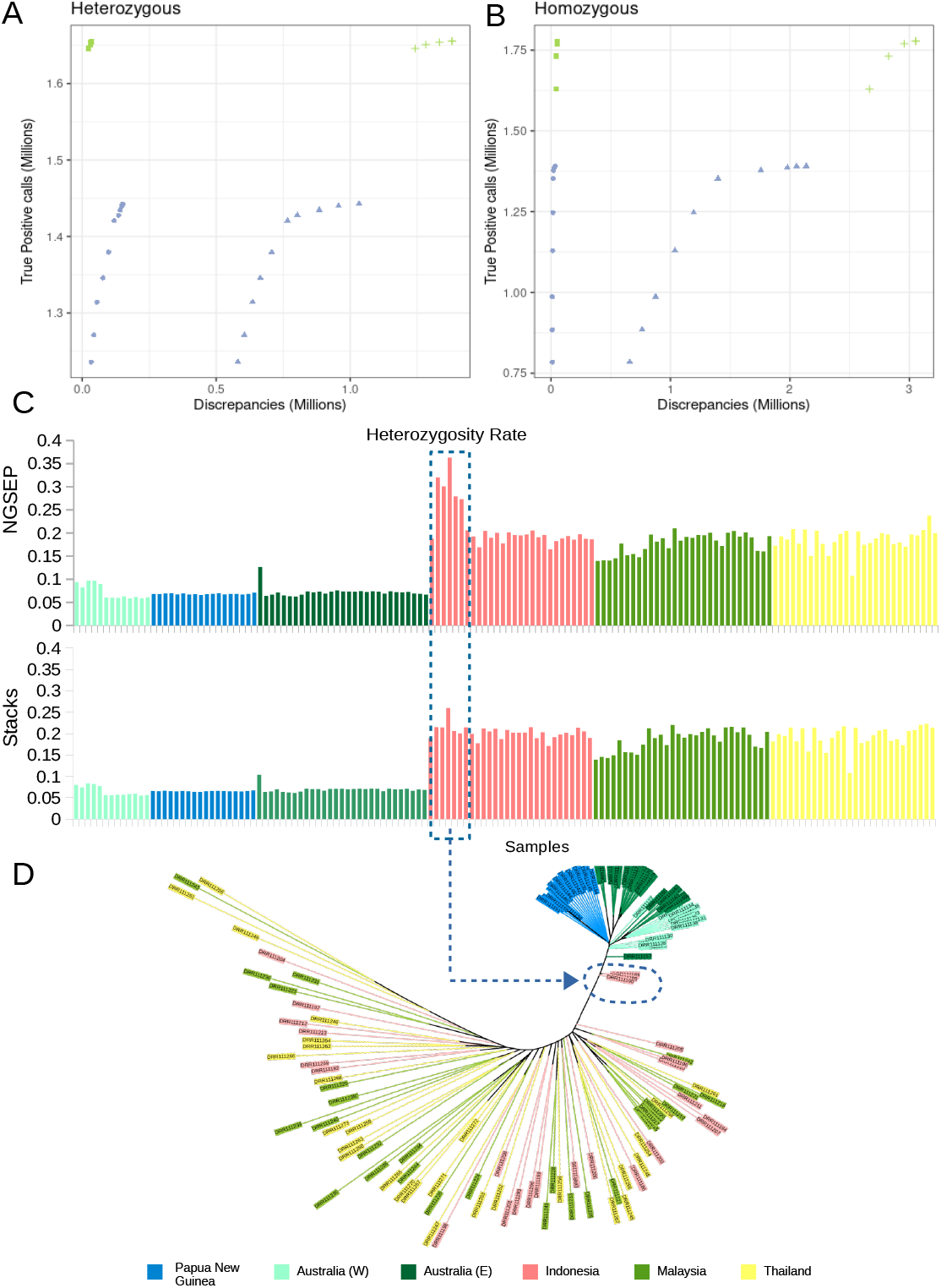
Sea bass diversity population. A. Number of matching non-reference genotype calls between de-novo and reference-based approaches as a function of two types of discrepancies: genotype differences within matching SNPs (solid lines) and non-reference genotype calls in SNPs identified only with the de-novo approach (dashed lines). C. Heterozygosity rate per individual calculated from genotype calls produced by NGSEP and Stacks. D. Dendrogram of genetic distances between the samples inferred from the genotype calls generated by the de-novo analysis of NGSEP.

Although the de-novo aproaches recovered in general less SNPs compared to the reference-based analysis, the distribution of diversity statistics was consistent for all strategies (Supplementary figure 3). This suggests that the SNPs and genotype calls obtained by the de-novo analysis of NGSEP can be used to investigate the diversity of the population. We built a dendrogram of genetic distances between samples inferred from the genotype calls obtained from the de-novo approach, filtering to keep only SNPs with at least 100 individuals genotyped with quality score greater than 40, and with a minor allele frequency greater than 0.01. Figure 3.D shows that samples are clustered according to geographical location, specifically the samples from Indonesia, East Australia and papua-New Guinea. This clustering is consistent with that reported by Wang and collaborators [36]. Population structure also seems consistent with the sample heterozygosity rates (Fig. 3.C), which clusters the samples in at least two groups: the population of Australia and Papua New Guinea, with heterozygosity rates of about 0.1 and the samples of Indonesia, Thailand and Malaysia with heterozygosity rates larger than 0.2. The dendograms obtained with both NGSEP (Fig. 3.C) and Stacks (Supplementary figure 4) show five samples that seem to be hybrids between the two major groups. However, heterozygosity rates for these samples reported by Stacks look similar to those for the samples from Indonesia, Thailand and Malaysia, whereas NGSEP reports higher heterozygosity rates for this samples (Fig.3.C samples inside the dashed blue rectangle).

### 2.5 Tetraploid Potato diversity population

Finally, to assess the accuracy of NGSEP de-novo on polyploid species, we analyzed SLAF-seq data from a tetraploid potato *(Solanum tuberosum)* biparental population of 100 individuals. As detailed in table 1, only GBS-SNP-CROP is able to perform polyploid variant genotyping. Unfortunately, barcodes required to operate this tool were not part of the available data for this population. Hence, we performed a comparison of genotype calls on shared variants considering only the reference based and the de-novo analysis performed by NGSEP. Figure 4 A shows that in this case the number of heterozygous genotype calls is much larger than that of homozygous genotype calls. Consistent with the results for the sea bass population, the percentage of discrepancies in heterozygous sites is overall larger than in homozygous sites (according to the reference-based analysis). Although the number of genotype differences is much larger than that observed in the sea bass population, it reduces to the same levels filtering by a minimum quality score of 60.

**Fig. 4:**
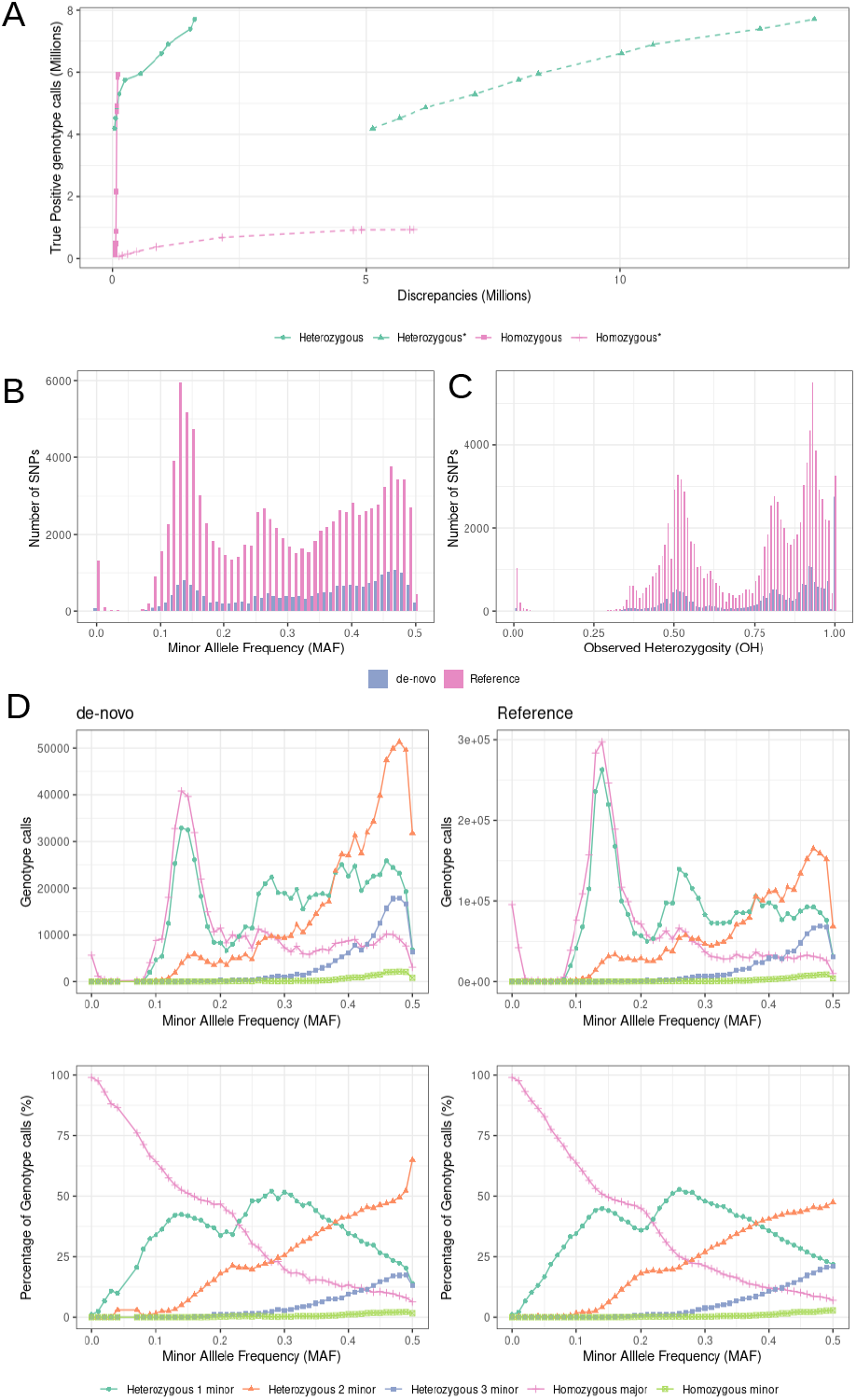
Tetraploid potato biparental population. A. Number of matching non-reference genotype calls between de-novo and reference-based approaches as a function of two types of discrepancies: genotype differences within matchin SNPs (solid lines) and non-reference genotype calls in SNPs identified only with the de-novo approach (dashed lines). B-C. Distribution of minor allele frequency (B) and observed heterozygosity (C) for the genotype calls produced by the de novo analysis and the reference-guided analysis of NGSEP on SNPs genotyped with quality 60 in at least 90 individuals. D. Number and percentage per MAF category of genotype calls discriminated by call type.

After filtering the different datasets keeping only variants with at least 90 individuals genotyped with genotype quality score larger than 60, the distribution of minor allele frequency (4 B) shows three peaks in 0.12, 0.25 and close to 0.5. Assuming one homozygous parent, these peaks correspond to cases in which the heterozygous parent has respectively one, two and four copies of the minor allele. The expected peak at 0.375 for a heterozygous parent with three copies of the minor allele is less evident, probably because one hundred individuals are not enough to differentiate the MAF distribution of SNPs with expected MAF around 0.5 and expected MAF around 0.375, also taking into account that three of the four cases with heterozygosity in both parents also have these expected MAF values. The clearest peak is observed for the reference-based analysis at 0.12, suggesting that homozygosity in one parent and heterozygosity with only one copy of the minor allele is the most frequent case between the parents. The distribution obtained by the de-novo analysis is consistent with that obtained from the reference-based analysis but the number of filtered SNPs (20,907) is much smaller than that of the reference-based analysis (104,797). The reduction in number of SNPs seems to be more pronounced for the category with MAF around 0.12. Conversely, less than 100 SNPs predicted by the de-novo analysis have MAF below 0.05, whereas this number is 1,475 for the reference-based analysis. The distribution of observed heterozygosity (4 C) does not show a peak around 0.25, which would be expected if the species were diploid but shows the first peak at 0.5, which is the expectation for SNPs with MAF around 0.12 in tetraploid species. Other peaks at 0.75 and 0.9 are also expected for SNPs with larger MAFs.

Considering that there are five possible types of genotype calls for biallelic variants in tetraploid species (two homozygous and three heterozygous), we calculated the number of genotype calls discriminated by call type and MAF. Figure 4 D shows that counts for both the reference-based analysis and the de-novo analysis are consistent with expectations. Homozygous calls for the major allele are predominant for SNPs with low MAF. For SNPs with MAF close to 0.12, calls are 50% homozygous for the major allele and 50% heterozygous with one copy of the minor allele. At MAF around 0.25, heterozygous call with one copy of the major allele become the most abundant keeping a percentage close to 50% whereas the homozygous calls reduce to 25% matching the heterozygous calls with two copies of the minor allele. The latter category grows to match the category with one copy of the minor allele at 0.37 and become the most abundant with 50% of the genotype calls for SNPs with MAF 0.5. Heterozygous calls with one copy of the minor allele reduce in percentage up to 20% at MAF 0.5, matching the calls with three copies of the minor allele. Finally homozygous calls for the major allele decrease to about 7% to get close to the homozygous calls for the minor allele, which grow to about 2% for SNPs with MAF close to 0.5.

### 2.6 Computational Performance

For benchmarking of computational efficiency, two main metrics were considered: run time and disk space. Disk space considers the amount of storage necessary to accommodate databases or supplementary files necessary for running the tool. Figure 5 shows the comparison of running times per Gb of input data between tools relative to the number of cores used by each tool. It evidences as well the fact that both run time and disk space usage increase linearly for the de-novo analysis implemented in NGSEP. Comparison between tools shows that our algorithm is faster than both PyRAD and Stacks for Gb of input data, and the reference guided analysis only comes close to it if the time required to aligned the reads to the reference genome is not taken into account. Even with one processor and less than 16Gb of RAM, our algorithm took 6 hours to process the rice population. For the sea bass population, NGSEP took 10 hours with 16 cores, whereas Stacks took 18 hours with 32 cores. Finally, NGSEP took 49 hours to process the potato population using 32 cores.

**Fig. 5:**
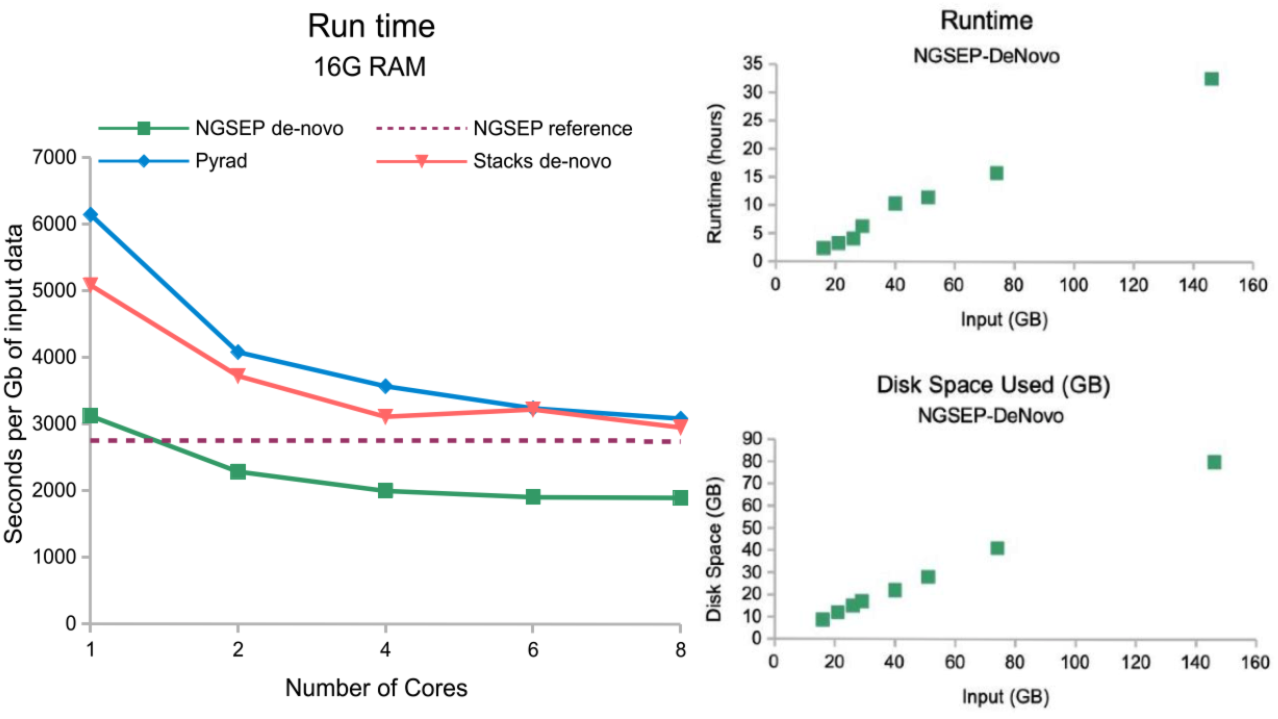
Experiments for run time were conducted in the High Performance Computing cluster of Universidad de Los Andes. RADProc was excluded from this analysis because it does not incorporate variant calling, which is an important part of the process. For the reference-guided analysis, the time required to align the reads to the reference genome was excluded.

## 3 Discussion

The use of high throughput sequencing (HTS) to perform cost-effective genotyping of thousands of markers across populations have become a tool of choice in population genomics. We developed an efficient algorithm to perform reference-free clustering of genotype-by-sequencing (GBS) reads and proved its effectiveness for different protocols, including double enzyme RAD sequencing (ddRAD) and SLAF-Seq, and for plant and animal species with different ploidies and heterozygosity rates. Our algorithm is able to cluster and process large amounts of GBS reads with similar accuracy and better efficiency, compared to widely used software tools. The voting strategy implemented for non-exact clustering of k-mers is both memory efficient (2 million clusters can be stored in 2Gbp of memory) and time efficient. Our algorithm was integrated in the Next Generation Sequencing Experience Platform (NGSEP), which facilitates the integration of a state-of-the-art algorithm for variants detection and genotyping [34]. This step is critical because some of the current tools implement well designed algorithms for clustering and consensus calling, but then implement simple variant calling models which in some cases are based only on read counts, ignoring the information contained in base quality scores [23]. Our results indicate that some of these methods tend to be over conservative, producing raw datasets with an inflated rate of missing data and low sensitivity once filtering is applied.

As evidenced by the analysis of the rice population, our algorithm retains a good sensitivity, even with datasets sequenced at low depths with a single-end protocol. The accuracy obtained with the de-novo algorithm ranked first among de-novo approaches and only ranked below the reference-based analysis. In particular, about 15% of the variants predicted by Stacks showed observed heterozygosity larger than 0.9, which is not expected for a population with six rounds of inbreeding. We also assessed the accuracy of our read clustering strategy on a sea bass population for which the ddRAD protocol was followed to sequence paired-end reads at high depth. Although for this dataset, Stacks showed better sensitivity than NGSEP, the percentage of homozygous genotype discrepancies between the results of Stacks and the reference-based analysis suggests that also in this case NGSEP achieves a better overall genotyping accuracy. Diversity analysis of the population resembles the structure reported in the original study, but NGSEP unveils five samples with a higher heterozygosity rate that also appears in the dendogram as a different group [36].

Finally, we assessed the accuracy of NGSEP to call variants on polyploids analyzing an F1 population of tetraploid potato [37]. The SLAF-Seq protocol was followed in this population to obtain paired-end reads at high depth [33]. The results obtained with this dataset indicate that the variant detection algorithm implemented in NGSEP has good accuracy to identify heterozygous and homozygous genotype calls. Even after genotype quality filters, the number of SNPs identified in this population exceeded 20,000 markers for the de-novo analysis. Compared to the analysis on diploid species, a more stringent filtering of genotype quality score is required to achieve comparable precision levels. This implies that, as expected, higher depth is required to achieve good accuracy in polyploid species. Once quality filters are applied, diversity statistics indicate that the SNPs genotyped with NGSEP show the segregation patterns expected for a tetraploid F1 population. Moreover, we verified that predicted allele dosages for heterozygous calls are consistent with the expected Mendelian inheritance that can be inferred from the calculated MAF of each SNP.

The experiments on the three datasets indicate that the clustering and genotyping algorithms presented in this manuscript achieve the best computational efficiency, particularly in runtime and disk usage. Particularly compared to Stacks, NGSEP is able to process similar datasets with two thirds of the CPU time. The cost of this improved efficiency seems to be a lower sensitivity in some datasets. Taking into account that GBS protocols are not meant to capture the complete set of SNPs segregating in a population, it could be argued that procedures to analyze GBS data should try to maximize genotyping accuracy preserving an adequate number of SNPs instead of maximizing sensitivity. However, as future work we will keep investigating on clustering alternatives that allow to increase sensitivity without losing genotyping accuracy and computational efficiency. It is worth mentioning that in general the de-novo analysis can be executed in shorter runtime compared to the reference-based analysis.

Achieving good computational efficiency was one of the main goals of this study, not only for benchmarking but also because we wanted to make it possible for researchers to run analysis of populations with hundreds of individuals with the computational resources of a desktop computer (about 8 cores with 16 GB of RAM and 1Tb of hard disk). To provide a good user experience we developed our algorithms as components of NGSEP to make it possible to execute the complete analysis on any operative system and with minimal installation requirements. The process requires standard fastq files as input and outputs a standard VCF file relative to a consensus sequences file in fasta format. Although this is also the case for other tools, NGSEP does not require any naming convention on filenames and especially on read ids. The clustering procedure performed by NGSEP also does not require reads to be of the same length, which allows to use a larger percentage of the raw sequencing data. Consensus sequences and SNPs can be mapped to a reference genome to provide genomic context to variants already identified by the de-novo analysis. Moreover, the de-novo analysis can be completely executed through the new graphical interface implemented in Java FX for version 4 of NGSEP (manuscript in preparation). Taking into account all these features, we expect that this new development will facilitate data analysis for a large number of researchers assessing genetic diversity within populations of a wide range of species.

## 4 Methods

### 4.1 Software development

The read clustering algorithm presented in this manuscript was implemented in Java 11 as part of the Next Generation Sequencing Experience Platform (NGSEP). This allows to take advantage of the object model previously implemented in NGSEP including internal representation and management of raw reads, alignments and variants. Moreover, the variants detector of NGSEP was integrated to discover and genotype variants on reads aligned to consensus sequences. Experiments included in this manuscript were performed with the released version 4.0.3 of NGSEP available as an open source product in sourceforge (http://ngsep.sf.net). Live development of NGSEP is also available in our github repositories (https://github.com/NGSEP).

### 4.2 Datasets used for benchmarking

We used three different datasets to assess the accuracy of the algorithm presented in this manuscript on different species and following different genotyping protocols 2. First, we reanalyzed publicly available single-end GBS data from an interspecific rice (*Oryza sativa)* biparental population of 176 recombinant inbred lines (IR64 x Azucena) [32]. Samples were sequenced using a 384-plex GBS protocol (140M, 94bp reads). The IRGSP1.0 reference genome [15] was used for read alignment in the reference-based approach and for mapping of consensus sequences and translation of variants in the de-novo approach.

**Table 2:** Datasets used for validation.

|  | Pop. description | Protocol | Dataset Size | Genome size |
| --- | --- | --- | --- | --- |
| Rice *O. Sativa* | Biparental inbred, diploid | GBS, single-end | 174 samples, 11Gb | 430Mb |
| Sea Bass *L. Calcarifer* | Diversity, diploid | dd-RAD, paired-end | 146 samples, 144Gb | 648Mb |
| Tetraploid Potato *S. Tuberosum* | Biparental, tetraploid | SLAF-Seq, paired-end | 100 samples, 1,500Gb | 840Mb |

As a second case for benchmarking, we reanalyzed Double Digest RAD (ddRAD) paired-end sequencing data from a diversity population of 146 sea bass *(Lates Calcarifer*) individuals [36]. This dataset was sequenced at a larger depth compared to the rice population, and it was useful to evaluate the potential of the algorithm to generate datasets useful to perform diversity analysis. We downloaded from NCBI the assembly released by [35] to use it as reference for read mapping, consensus sequences mapping and translation of de-novo SNPs.

Finally, we reanalyzed SLAF-Seq paired-end data from 100 individuals of an F1 biparental population of tetraploid potato *Solanum tuberosum*, generated to build a genetic map for this species [37]. Similar to the case of the rice population, this controlled population allows to test expectations on the distribution of population statistics. A recently released chromosome level genome of diploid potato was used as reference [26].

### 4.3 Tools used for comparison

We selected tools currently available and used for GBS de-novo analysis. All tools were run using default parameters unless a change was necessary for correct functioning. Filtering of VCFs was done using the VCFFilter capability of NGSEP [34].

Stacks was installed by downloading the package available at https://catchenlab.life.illinois.edu/stacks/. Given that stacks requires all reads to be the same length, after cleaning the datasets of adaptor contamination, reads were filtered and trimmed to preserve the same length. For the rice population, reads longer than 70bp were trimmed to this length and shorter reads were filtered out. For the Seabass population, only reads without adapter contamination (about 75% of the total) were retained. The scripts *denovo_map.pl* and *ref_map.pl* were used to run the de-novo analysis and the reference-based analysis respectively (Supplementary table 1). Consensus sequences generated by Stacks were used to translate the VCF to the coordinates of the reference genome.

RADProc was built from source from its GitHub repository https://github.com/beiko-lab/RADProc. Given that it is a tool that exclusively seeks to improve Stacks performance when building the catalogue, it does not perform variant calling. To perform variant calling, the catalogue created by RADProc was passed to sstacks, tsv2bam, gstacks and populations modules from Stacks 2.2 to produce a VCF. The catalogue created by RADProc describing the consensus sequences, was used to translate and validate the variants found.

PyRAD was built from source from its GitHub repository https://github.com/dereneaton/pyrad/releases. Python modules Numpy and SciPy were installed, as well as MUSCLE and VSEARCH [8, 28] which are required for clustering. PyRAD parameters are set from a configuration file. Mindepth, maxN and clustering threshold were all set to the suggested values (5, 4, 85% and 90%). Unfortunatey, PyRAD does not output a consensus file and therefore it was not possible to translate and validate individual variants.

GBS-SNP-CROP is a pipeline that runs seven PERL scripts in sequence. Unfortunately, GBS-SNP-CROP requires a barcodes file, even if the data has already been de-multiplexed, which was not available for the tested populations. For this reason this tool could not be tested.

### 4.4 Accuracy measures

Although the goal of the de-novo analysis of GBS data is to provide information of genetic diversity in species without a reference genome, we chose the datasets presented in this study on species having a reference genome, to be able to compare in all cases variants generated de-novo with variants generated by a reference-based analysis. Both the reference-based analysis ([34]) and the de-novo analysis (this manuscript) implemented in NGSEP were executed for each benchmark dataset (see detailed parameters in the supplementary text). The reference-based analysis of Stacks was also executed in the rice population but the calls of the de-novo analysis of both Stacks and NGSEP showed better coincidence with the reference-based analysis of NGSEP. Hence, for all datasets we considered the variants and genotype calls generated by the reference-based analysis of NGSEP as gold-standard for comparison. Variants generated by each de-novo pipeline were translated to reference coordinates using the command *VCF Relative Coordinates Translator* of NGSEP. In brief, this functionality first aligns the consensus of each cluster to the reference genome and then uses the alignment to map the reported variant positions relative to the cluster consensus back to variant positions relative to the reference.

Initially, we tried to assess accuracy by coincide of SNP locations, after translation of de-novo variants to the reference genome. However, the accumulation of SNPs called out of only one erroneous non reference genotype call generate an inflated estimation of false positives for all tools and does not really allow to evaluate accuracy of genotype calls. Hence, as a measure of sensitivity, we calculated the the total number of matching non-homozygous reference genotype calls within SNPs identified by both the de-novo approach and the reference-based approach. This was constrasted with the number of discrepancies which were classified in two types: 1) non-matching genotype calls within SNPs identified by both the de-novo approach and the reference-based approach, and 2) non homozygous reference genotype calls within SNPs only identified by the de-novo approach. This approach is implemented in the script *ngsep.benchmark. Genotype Based Population SNP Gold Standard Comparator* available with the distribution of NGSEP.

For the rice population, reference-free benchmarking was performed measuring sensitivity as the number of variants that follow the expected segregation pattern (MAF ? 0.1 and OH ≤ 0.1) for the population and contrasting this number with the number of genotype calls that should be erroneous. Although theoretically an F6 population should still an observed heterozygosity around 2%, we considered all heterozygous genotype calls for this population as possible errors. Moreover, homozygous calls to the minor allele within SNPs having low MAF are also considered erroneous. This procedure is implemented in the script *ngsep.benchmark.Quality Statistics Inbred Biparental Families* available with the distribution of NGSEP.

Finally, for the tetraploid potato F1 biparental population, the number of genotype calls was counted discriminating each of the five possible genotype calls in a biallelic variant, which correspond to the number of copies of the minor allele within the individual (from zero to four). Counts were also discriminated by MAF category from zero to 0.5 in intervals of 0.01. Counts were plotted having the MAF category in the x-axis to assess visually if the counts are percentages were consistent with expectations according to the type of population. Counts are calculated by the script *ngsep.benchmark. Quality Statistics Tetraploid F1 Families* available with the distribution of NGSEP.

## 5 Acknowledgements

We are thankful for I.Ceron-Souza insights while conceiving this project.

