## Supplementary for "Robust and efficient software for reference-free genomic diversity analysis of GBS data on diploid and polyploid species"

### Supplementary tables

Supplementary table 1: Parameters used to run the different software tools on each benchmark dataset.

| Population | Software | Parameters |
|---|---|---|
| Rice biparental F6 | NGSEP de novo | java -XX:+UseSerialGC -Xmx16g -jar NGSEPcore_4.0.3.jar DeNovoGBS -i readsGBSRice -d sampleIdsFiles.txt -t 4 -maxBaseQS 30 -h 0.0001 -minQuality 40 -o GBSRice_NGSEPDeNovo403 |
| | Stacks de novo | stacks-2.54/scripts/denovo_map.pl --samples readsL70 --popmap popmap.txt -o StacksDeNovo -M 3 -n 2 -X "ustacks: -m 2" -T 8<br>stacks-2.54/populations -P StacksDeNovo -M popmap.txt --vcf -t 8 |
| | NGSEP reference | java -XX:+UseSerialGC -Xmx6g -jar NGSEPcore_4.0.3.jar MultisampleVariantsDetector -maxAlnsPerStartPos 100 -maxBaseQS 30 -minMQ 40 -knownSTRs nipponbare_TIGR7_trf_2.7.7.80.10.20.50_filter.txt -r nipponbare_TIGR7.fa -o GBSRice_NGSEPref403.vcf mapping/*bam |
| | Stacks reference | stacks-2.54/scripts/ref_map.pl --samples mappingL70 --popmap popmap.txt -o StacksMapping -T 8 -X "populations: --vcf" |
| | RADproc | RADProc -t gzfastq -f readsL70 -o RADproc -M 2 -m 3 -n 2 |
| | PyRAD | pyrad_v.3.0/pyRAD -p params.txt -s 234567 |
| Seabass diversity | NGSEP de novo | java -XX:+UseSerialGC -Xmx32g -jar NGSEPcore_4.0.3.jar DeNovoGBS -c 8000000 -t 16 -maxBaseQS 30 -i trimmed5bpAll -d sampleFiles.txt -o Seabass_5bpAll_NGSEP403 |
| | Stacks de novo | stacks-2.54/scripts/denovo_map.pl --samples trimmedL146 --popmap popmapFinal.txt -o stacks --paired -M 3 -n 2 -X "ustacks: -m 5" -T 32<br>stacks-2.54/scripts/populations -P stacks -M popmapFinal.txt -t 16 --vcf |
| | NGSEP reference | java -XX:+UseSerialGC -Xmx16g -jar NGSEPcore_4.0.3.jar MultisampleVariantsDetector -maxAlnsPerStartPos 100 -maxBaseQS 30 -minMQ 40 -knownSTRs nipponbare_TIGR7_trf_2.7.7.80.10.20.50_filter.txt -r nipponbare_TIGR7.fa -o GBSRice_NGSEPref403.vcf mapping/*bam |
| Potato biparental F1 | NGSEP de novo | java -XX:+UseSerialGC -Xmx32g -jar NGSEPcore_4.0.3.jar DeNovoGBS -c 8000000 -t 32 -maxBaseQS 30 -ignore5 5 -ignore3 5 -i potatoReads -d sampleFiles.txt -o PotatoBiparental_NGSEPdeNovo403 |
| | NGSEP reference | java -XX:+UseSerialGC -Xmx16g -jar NGSEPcore_4.0.3.jar MultisampleVariantsDetector -maxAlnsPerStartPos 100 -maxBaseQS 30 -minMQ 20 -r potato_genome_assembly.v6.1.fa -o PotatoBiparental_NGSEPref403.vcf mapping/*bam |

### Supplementary figures

Supplementary figure 1. Distribution of quality scores generated by different software tools on the rice biparental population

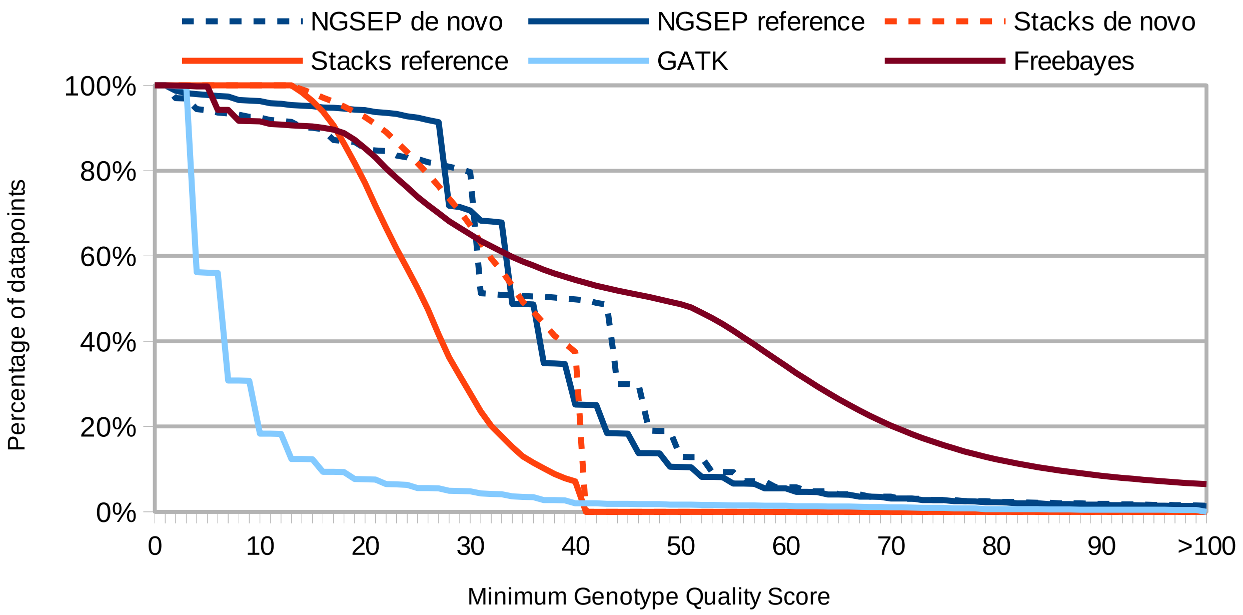

Supplementary figure 2. Density of genotype calls for SNPs discovered in the sea bass diversity population by the de-novo pipelines of NGSEP and Stacks

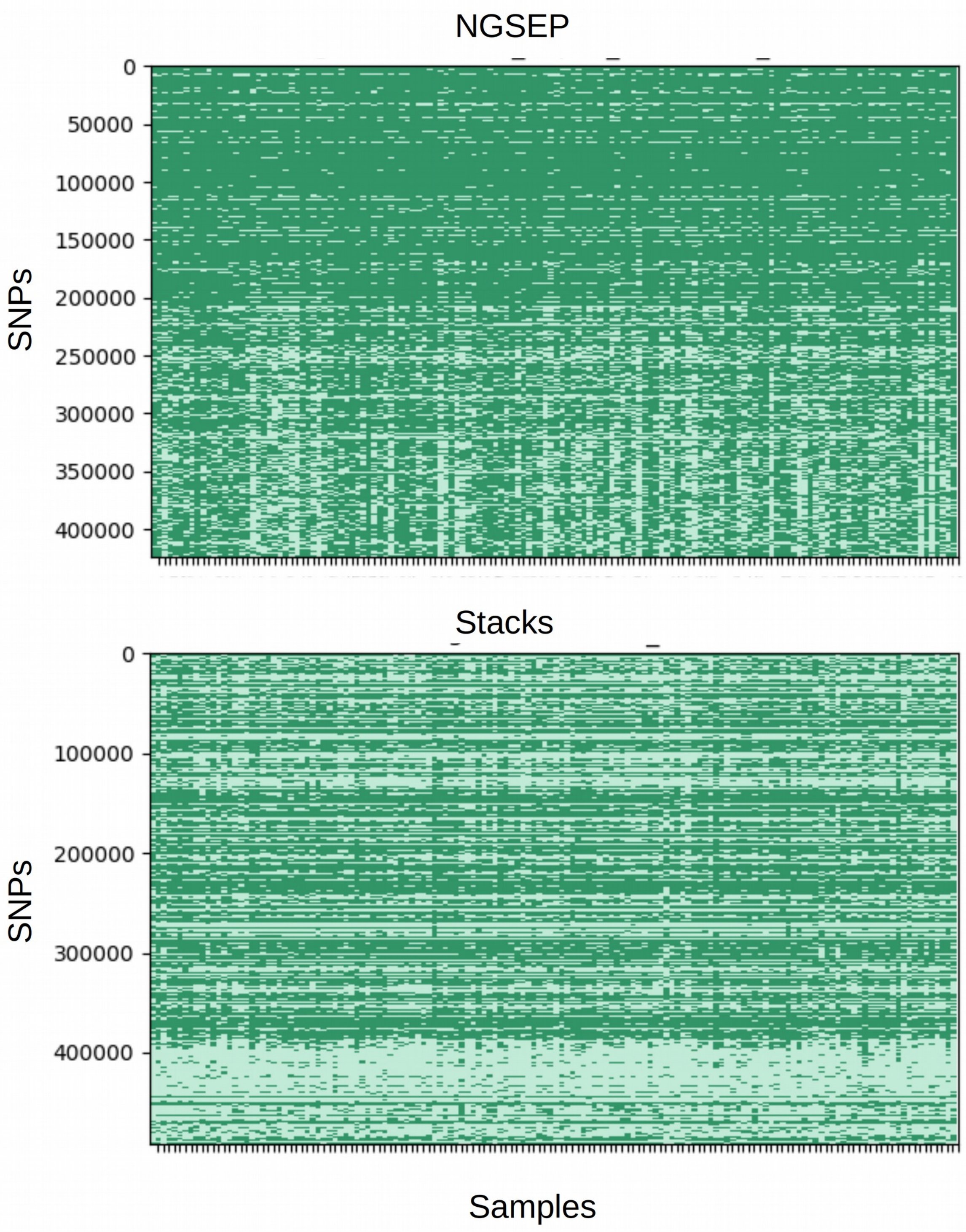

Supplementary figure 3. Distribution of minor allele frequency and observed heterozygosity for the genotype calls produced by the de novo analysis of NGSEP and Stacks and the reference-guided analysis on SNPs genotyped with quality 40 in at least 100 individuals

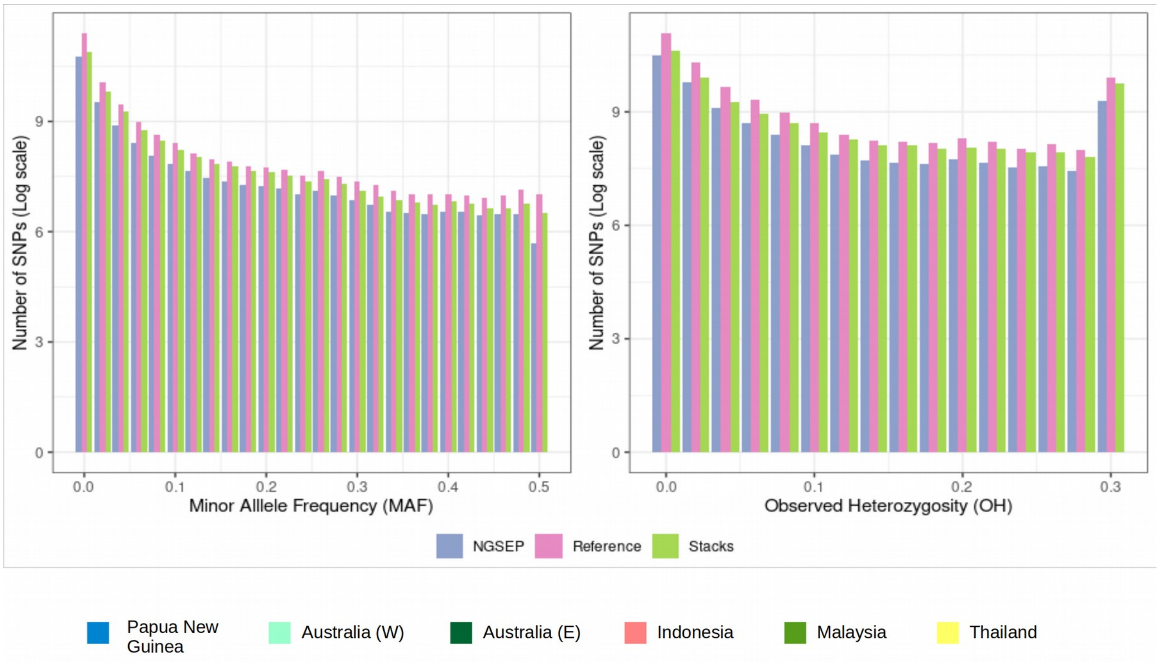

Supplementary figure 4. Dendrogram of genetic distances between the samples inferred from the genotype calls generated by the de-novo analysis of Stacks.

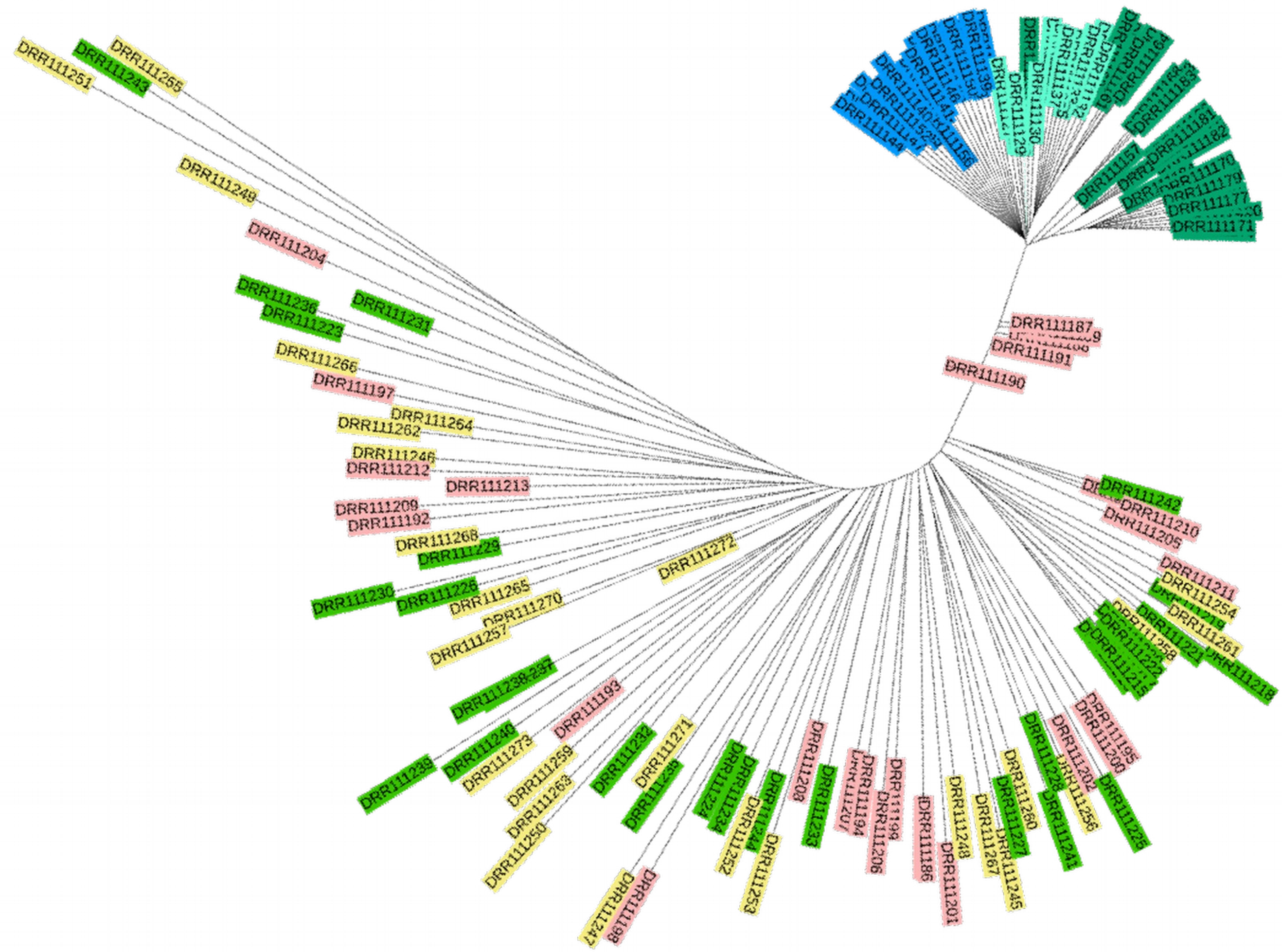
